# Diploid gametes in maize by mutation of A-Type cyclins: a step towards apomeiosis and synthetic apomixis

**DOI:** 10.1101/2025.05.16.654085

**Authors:** D.J. Skinner, A.H. Gaikwad, J.A. Fenner, J. Green, T. Kelliher, M-J. Cho, V. Sundaresan

## Abstract

Hybrid crops are agriculturally desirable due to heterosis but are costly or difficult to produce. Synthetic apomixis, clonal reproduction through seed, offers the ability to fix hybrid vigor. Two components are needed to achieve this goal: the formation of diploid gametes identical to the maternal parent through apomeiosis, and the induction of embryogenesis in the egg cell without fertilization, known as parthenogenesis. In maize, parthenogenesis was achieved by egg cell expression of the transcription factor *ZmBABY BOOM 1* but a viable apomeiosis strategy has not been reported. In the MiMe (Mitosis instead of Meiosis) system, in addition to mutants that skip recombination and sister chromatid adhesion, mutation of genes involved in cell cycle control during meiosis is needed to skip the second division and ensure diploid gametes. In this report we describe the effect of mutation of maize A-type cyclin genes with similarity to Arabidopsis *TARDY ASYNCHRONOUS MEIOSIS* (*TAM*). In double mutant plants, we find that diploid gametes are formed with high efficiency and that the progeny are tetraploid. These genes provide a viable route towards creating synthetic apomixis in maize.

## Introduction

The large-scale use of hybrid seeds over the past century has contributed to a revolution in agriculture. Hybrids produce significantly higher yields but due to genetic segregation, these yields are not maintained by seed propagation. In maize and other major crops, hybrid seeds are newly generated each planting season by cross-pollination between inbred parents. Commercial hybrid maize production uses large scale emasculation of female parents, requiring the use of fossil fuel resources and extra land for the growth of male parents. The economic and environmental costs would be significantly reduced if hybrid plants can be made to reproduce asexually without genetic segregation, as occurs naturally in apomixis.

Apomixis, asexual reproduction through seeds, has evolved in many angiosperm lineages (Hand and Koltunow, 2014) and is found in several grass species. Recently, successful clonally reproducing rice hybrids via synthetic apomixis has been achieved (Khanday et al., 2019; Vernet et al., 2022). The two key aspects of synthetic apomixis are parthenogenesis, embryo growth without fertilization, and apomeiosis, the substitution of meiosis with mitosis (called MiMe). MiMe produces diploid clonal gametes by knockouts of three meiotic processes (d’Erfurth et al., 2009; Mieulet et al., 2016). In rice and maize, parthenogenesis has been achieved by ectopic egg cell of expression of the transcription factor BBM (BABY BOOM), an embryogenic trigger expressed in wild-type zygotes (Khanday et al., 2019; Skinner et al., 2023).

The *MiMe* (Mitosis instead of Meiosis) phenotype arises from combining mutants that do not undergo the second meiotic division with those lacking meiotic recombination (e.g., *spo11-1*) and sister chromatid cohesion (e.g., *rec8*). First shown in *Arabidopsis, spo11 rec8 osd1* triple mutants produce unrecombined, diploid gametes that self-fertilize to form tetraploid offspring (d’Erfurth *et al*., 2009). Similarly, rice *pair1 rec8 osd1* triple mutants also produce non-recombined diploid male and female gametes and this combination was used for synthetic apomixis (Mieulet *et al*., 2016). *spo11* and *pair1* mutants do not initiate double strand breaks, leading to absence of recombination, which is required for clonal gametes. *REC8* is a highly conserved protein that maintains cohesion of sister chromatids, and mutants of *rec8* in the MiMe context allow for separation of chromatids after meiosis I. Maize *SPO11-1* and *AFD1* have been characterized and show conserved function with the rice and Arabidopsis genes.

The third step to achieve viable unreduced gametes by MiMe requires exit from meiosis prior to the second division. For both mitosis and meiosis, exit at anaphase occurs when the activity of cell cycle components such as cyclins and cyclin-dependent kinases (CDK) cease. In meiosis, CDK1;A activity rises during meiosis I, then drops to allow separation of homologous chromosomes at anaphase I. This decrease in activity is caused by degradation of cyclins by the anaphase promoting complex/cylosome (APC/C), an E3 ubiquitin ligase which targets cell cycle components. Unique to meiosis, a second division (without DNA synthesis) occurs because the cyclin levels are not completely abolished. CDK activity increases again during meiosis II until anaphase II when complete inactivation by the APC/C allows sister chromatids to segregate to opposite poles. APC/C activity must therefore be tightly regulated to both ensure a second division and block further divisions after meiosis II, and this regulation is one point at which the cell cycle can be manipulated for an early meiosis I exit.

The regulation and function of the APC/C is complex and largely unknown in plants. OMISSION OF SECOND DIVISION1 (OSD1), encoding an APC/C inhibitor, is required for meiosis II entry in both Arabidopsis and rice. *osd1* mutants have been used to skip the second division in MiMe, a phenotype likely caused by higher activity of the APC/C during meiosis I. In tomato and many other species, the use of OSD1mutants for generating diploid gametes has been restricted, due to both mitotic and meiotic cell cycle functions being supplied by a single OSD1 gene, leading to lethal knockouts. In maize, while there are three putative OSD1 homologs, their expression suggests they may all be active in mitosis and therefore not useful targets for MiMe.

As an alternative, a meiotic A-type cyclin which may interact with CDKA;1, TARDY ASYNCHRONOUS MEIOSIS (TAM), is active during Meiosis I and essential for the entry to meiosis II in Arabidopsis (d’Erfurth et al., 2010)(Bulankova et al., 2010). *tam* mutants combined with Arabidopsis *spo11* and *rec8* mutants resulted in an alternative path to clonal diploid gametes, known as MiMe2. Similarly, *Sltam* mutants of tomato give rise to diploid gametes through MiMe2 when combined with *Slspo11* and *Slrec8* mutants (Wang et al., 2024). In this work, maize A-type cyclins that may have a similar function to Arabidopsis TAM were identified and tested for their ability to give rise to diploid gametes. This system could be used for apomeiosis, an essential step towards achieving synthetic apomixis in maize, which would be of significant agricultural benefit.

## Results

Maize CyclinA1;1 (Zm00001eb128470, *CYC2*) and CyclinA1;2 (Zm00001eb339350, *CYC6*) are A-type cyclins with high similarity to Arabidopsis *TAM*. BLAST searches of the maize reference genome (B73v5) were done to find any additional *ZmTAM* candidates, which were cross checked against a single cell male meiosis transcription data set (Nelms and Walbot, 2022). Candidate genes for cyclins controlling entry into Meiosis II are expected to be expressed preferentially in Meiosis I (Bulankova et al., 2013, 2010). Four cyclins with greater than 50% aa similarity to AtTAM in the cyclin domains also show Meiosis I expression. *CYC2* and *CYC27* (Zm00001eb350330) show declining expression as Meiosis II proceeds, while *CYC6* and CYC9 have continued expression during the gametophytic Pollen Mitosis I (Fig. 1). *CYC27* has very low to zero expression values in vegetative tissues, while *CYC2, CYC6* and *CYC9* have more widespread, higher expression values (Walley et al., 2016). Of these four genes, *CYC9* does not show significant amino acid similarity with *AtTAM* along the N-terminal region of the protein (Fig. S1). On the basis of amino acid similarity and expression, *CYC2, CYC6* and *CYC27* were chosen for further analysis.

**Figure 1.**
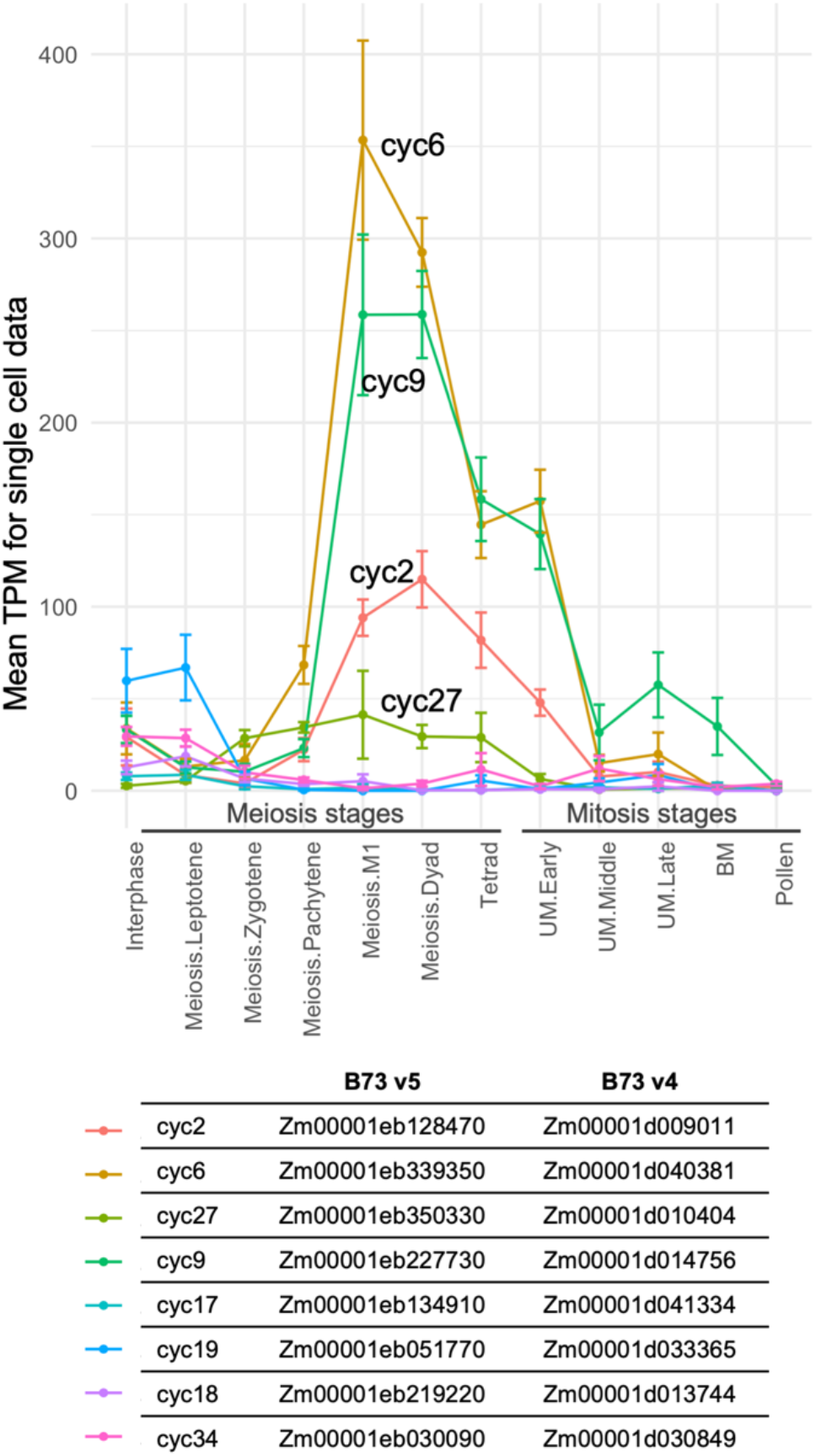
Transcription levels of select cyclin genes with similarity to *AtTAM* during male gamete development in maize. Data from single cell analysis of pollen development (Nelms, 2022).

We generated CRISPR-Cas9 constructs to target these genes singly and using multiplex editing in the B104 maize genotype. We checked for DNA sequence conservation between B73 and B104 and found that the coding regions shared only a few sequence variants, although the *CYC27* gene in B104 was not fully annotated. Two guide RNAs per gene were designed to target the CYC genes uniquely, despite similar sequence between *CYC2* and *CYC6*. For single gene targeting, homozygous knockout mutants were obtained in the T0 or T1 from at least 2 independent transgenic events for each gene (Table 1). For multiplex targeting, a tRNA-gRNA array was used with the same pair of guides for each gene, arrayed in two sets of three guides, each driven by their own OsU6 promoter. From this construct, five independent events gave rise to various combinations of gene knockouts in the T0, giving rise to double mutants in later generations.

**Table 1.**
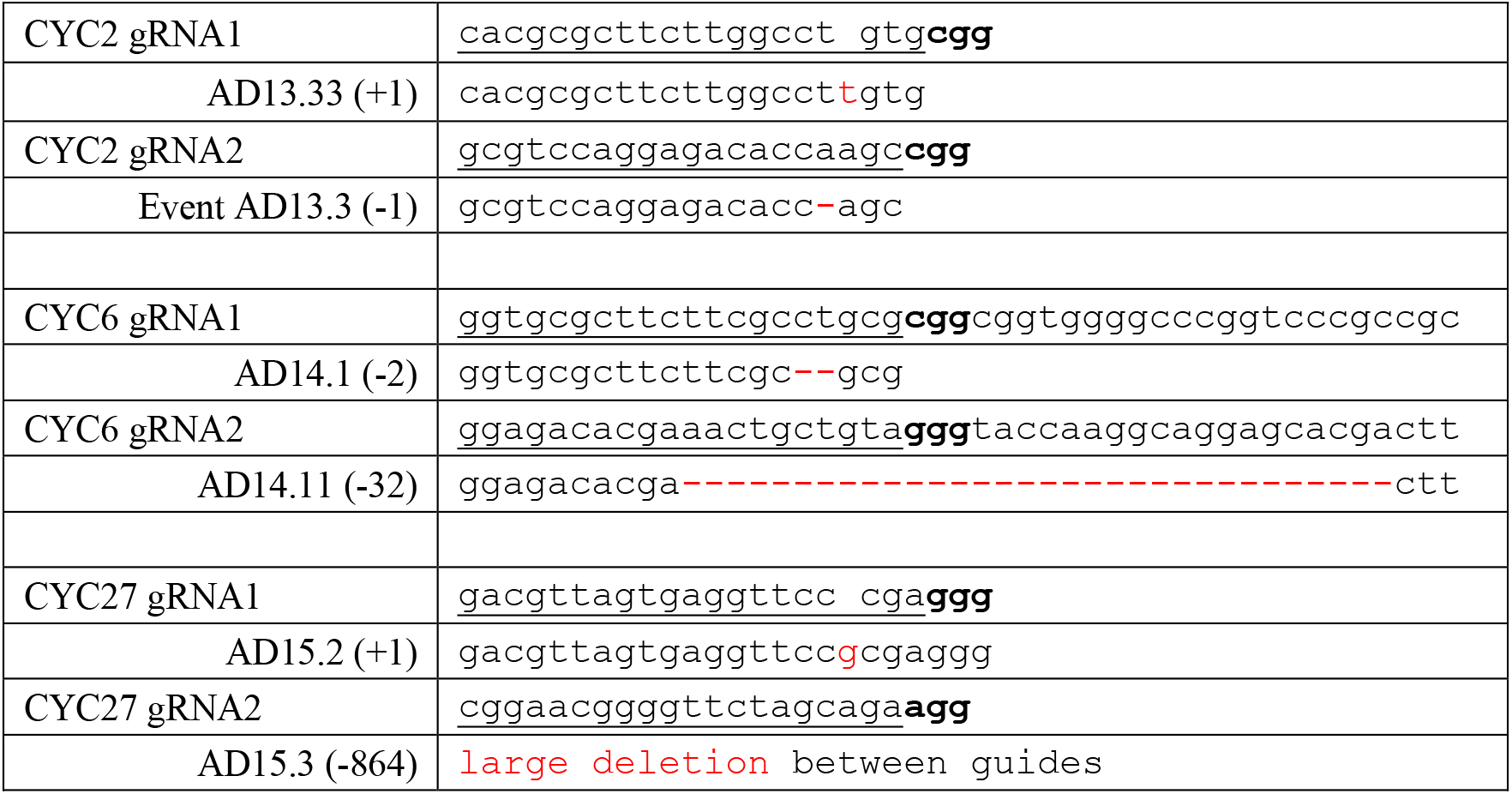
Guide RNA sequences for CYC genes (underlined) with PAM (bold) within regions being targeted. Representative edited sequences for single mutants from different independent transgenic events are shown in red.

If *cyc* mutants cause a disruption of meiosis II, the resulting phenotype would be diploid gametes. We assessed this phenotype by flow cytometry of nuclei collected after manual disruption of pollen (Fig.2). Single mutants showed normal haploid pollen, and progeny were diploid, showing that neither male nor female meiosis was disrupted to a detectable level. *cyc2 cyc6* double mutants from two events had reduced pollen quantity, as observed by narrower anthers and less pollen shed. The pollen from these mutants was haploid, as observed by flow cytometry. In contrast, *cyc2 cyc27* double mutants showed evidence of diploid pollen as small diploid peaks were observed in flow cytometry. In all preparations of double mutant individuals, nuclei appeared significantly degraded, or of reduced fluorescence, even though control samples prepared at the same time gave clear peaks. This meant that the ratio of haploid:diploid pollen could not be estimated from flow cytometry of pollen grains alone. Nevertheless, these results suggested that cyc2 cyc27 mutants were capable of producing diploid gametes.

**Figure 2.**
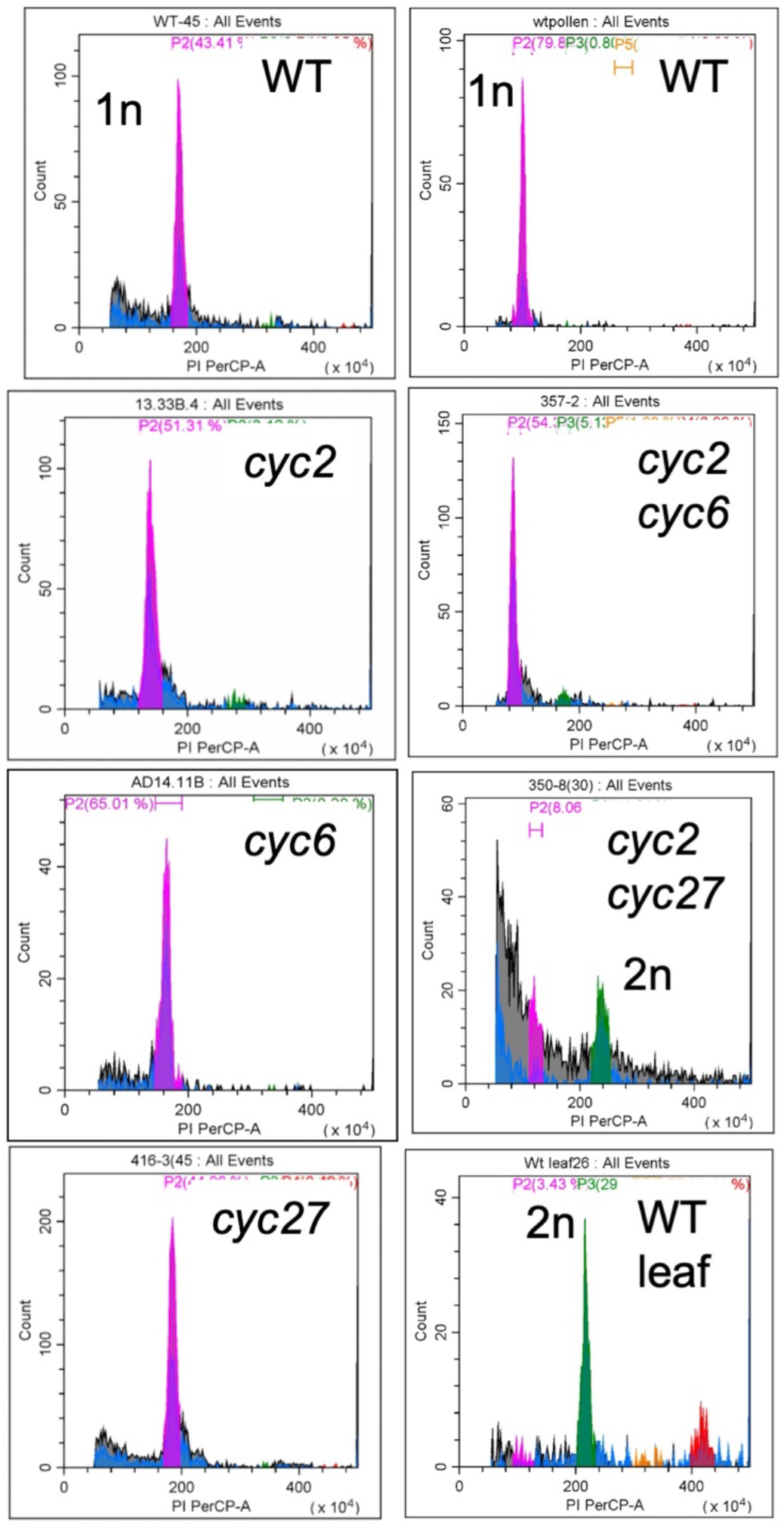
Flow cytometry of pollen and leaf nuclei from the wild type (WT) B104 controls and single and double mutants. 1n peaks are pink and 2n peaks are green. The WT leaf sample contains both 2n and 4n (red) peaks due to endoreduplication in some leaf cells.

Successful use of *cyc* mutants for synthetic apomixis would require female gametes to be diploid. Ploidy of female gametes is difficult to measure directly, but can be inferred by determining the ploidy of progeny from selfing. One T1 plant from event AD16.11 carried a homozygous -32 bp deletion in *cyc2* and was heterozygous for a +1 bp insertion in *cyc27* with no remaining Cas9 transgene. When selfed, the resulting T2 generation segregated several double mutant plants which showed diploid pollen. When these plants were selfed, the plants showed moderate fertility (roughly 50% of kernels appeared fertilized). From the fertilized kernels, some seeds aborted before maturity, but there was high percentage of filled kernels (74-92%) as compared with control *cyc2 cyc27/+* selfed ears (98% filled kernels, Fig 3d, g).

**Figure 3.**
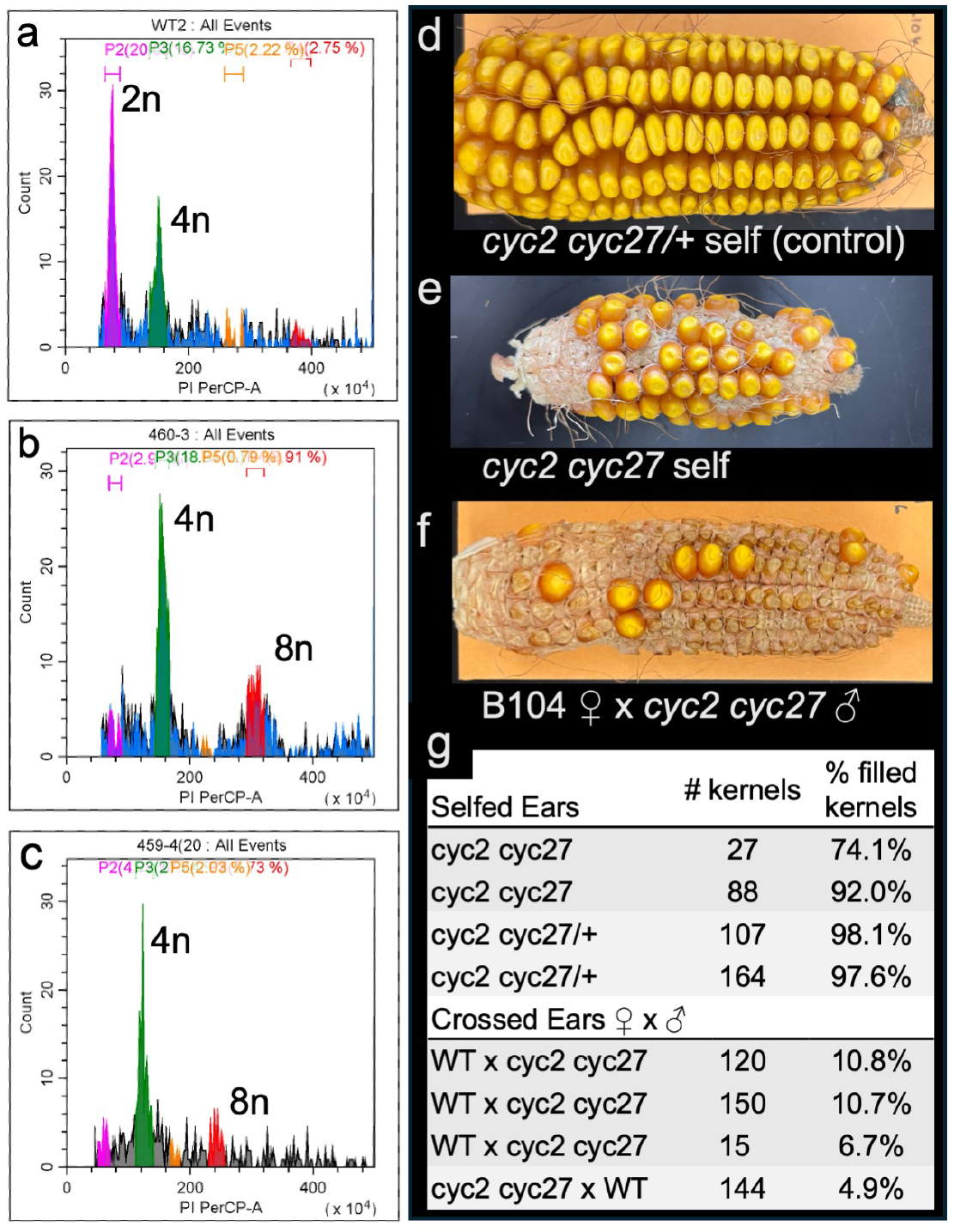
Analysis of progeny of *cyc2 cyc27* double mutants. a-c: Flow cytometry of wild type (WT) leaf (a) compared with two representative tetraploid progeny from selfs of two independently isolated *cyc2 cy27* mutants (b, c). WT leaf contains mostly 2n nuclei, with 4n nuclei from endoreduplication. The tetraploid progeny show mostly 4n nuclei with additional 8n nuclei as expected. (d-f): Ears arising from the self or crosses annotated in the figure. (d) Selfed ear from a *cyc2 cyc27/+* plant showing normal seed set. (e) Selfed ear from a *cyc2 cyc27* plant showing several mature, filled kernels but reduced fertility as many ovules were not fertilized. (f) WT ear pollinated with *cyc2 cyc27* pollen, showing that most kernels were fertilized but kernels did not develop properly (shriveled seeds). (g) The percentage of filled, mature kernels that developed on ears that were selfed or crossed showing that few kernels are formed when the gamete ploidy is not matched.

T3 progeny that germinated were tested by flow cytometry to determine ploidy. 24 T3 plants were tested and all were tetraploid (Fig. 3b). A second event (AD16.21) gave rise to a T1 plant that was homozygous *cyc2* (+1/+1 bp) *cyc27* (-1/-1 bp). This plant had diploid pollen and also gave rise to tetraploid plants (4 of 4 tested, Fig. 3c). This result shows that female gametes are also affected in Meiosis II and have diploid gametes. Unfilled or underdeveloped kernels, caused by failure of endosperm or embryo development, could be attributed to mitotic effects of the mutants, or to incomplete penetrance of the diploid gamete phenotype. If some haploid gametes are produced and fertilize a diploid gamete, the resulting unbalanced endosperm may not develop.

In order to determine the penetrance of the diploid gamete phenotype, reciprocal crosses were made between *cyc2 cyc27* and WT B104 plants. With B104 as the female parent, for seeds fertilized with diploid mutant pollen, the embryo would be triploid, while the endosperm would have a 2n:2n female to male ratio, which leads to failure to develop and kernel arrest in most cases (Sarkar, 1974). Therefore, any seeds that develop from this cross are most likely to have been the result of fertilization with haploid pollen. From three ears, we observed most fertilized kernels abort as expected, with just 6-11 % of kernels being normal sized (Fig 3e, g). This suggests that diploid pollen constitutes the majority of the functional pollen made by the *cyc2 cyc27* mutant. In the reverse cross, with *cyc2 cyc27* as the female, the endosperm would have a 4n:1n ratio, which is tolerated slightly more often for kernel development. Even in these crosses, we observed very few mature kernels (Fig 3f, g) implying that the penetrance during female meiosis is also high.

## Discussion

The production of clonal diploid gametes in maize, in combination with parthenogenesis or factors such as *ZmBBM* (Skinner et al., 2023) or haploid induction mechanisms such *ZmMTL* (Kelliher et al., 2017) would allow for synthetic apomixis in this crop plant. Few meiotic mutants have been described in maize that have diploid gametes, and most result in high male and female sterility, and low frequency of viable unreduced gametes. The *argonaute104* mutant produces a larger proportion of diploid gametes due to defects in small RNA processing, but shows extensive pleiotropy (Singh et al., 2011). Similarly, the *elongate1* mutant of maize produces diploid gametes, but the identity of the underlaying mutation is unknown, and 2n gametes are made in proportions too low for applied use (Barrell and Grossniklaus, 2005; Nel, 1975). The results of double mutation of *cyc2* and *cyc27* show that high proportions of diploid gametes are produced in largely phenotypically normal plants which has not been previously observed.

Altering components of cell cycle regulation has been very effective in creating diploid gametes, as observed in the Arabidopsis and rice *osd1* mutants when combined with *rec8* and either *spo11* or *pair1*. However, barriers to the use of *osd1* mutants include those crops such as tomato (Wang et al., 2024) and watermelon, (Pang et al., 2025) where mutation is lethal or deleterious due to defective mitosis and pleiotropic effects. As an alternative, *tam* mutants have also shown high proportions of diploid gametes in Arabidopsis and tomato. In these plants, *tam* in *MiMe2* leads to >85% tetraploid progeny, with the remaining progeny showing possible aneuploidy. In both plant types, pollen development lead to 2n gametes as well as a low proportion of 1n gametes and unbalanced products (aneuploids). In the maize *cyc2 cyc27* mutants, we likely also observed aberrant male gametes, which appear as nuclei or nuclear debris with less than 2n DNA content in flow cytometry. However, as in the tomato experiments, tetraploid plants are mainly formed in the progeny, showing that these mutants are viable for use in synthetic apomixis.

Synthetic apomixis in corn would affect both the production and availability of hybrid seeds worldwide. Maize is the main source of calories for many in the developing world, and many small-scale and subsistence farmers use lower-yielding, open-pollinated varieties as they cannot access the most improved hybrid seed. The results of this study demonstrate the feasibility of achieving apomeiosis, and consequently synthetic apomixis in maize.

## Methods

Guide RNAs for the CYC genes were chosen using the CRISPOR website and are shown in Table 1. The pair of guide RNAs for single gene targeting were cloned into an ENTRY vector pENTR-sgRNA4 which contains two different OsU6 promoters (Zhou et al., 2014). This vector was also used to express the multiplex tRNA-gRNA array. These guide RNA cassettes were recombined into a binary vector based on pZY101 carrying a maize-optimized Cas9 gene driven by the ZmUbi-ZmAdh1intron promoter and bialaphos resistance for selection in maize transformation. Transformations into the B104 genotype were performed at the Innovative Genomics Institute. T0 events were split into 1-5 plantlets and genotyped for presence of the transgene, and for edits in target genes.

Nuclei extraction from pollen was performed by bursting the pollen against a 40 uM filter in extraction buffer (Kron and Husband, 2012), followed by staining with propidium iodide. Leaf nuclei extraction was performed by chopping young leaves (Galbraith and Lambert, 2012). Flow cytometry was performed on a Beckman Coulter Cytoflex Flow Cytometer with standard protocols.

