## Supplemental Figure 1 for "Diploid gametes in maize by mutation of A-Type cyclins: a step towards apomeiosis and synthetic apomixis"

Supplemental Data

Figure S1 CLUSTAL O (1.2.4) multiple sequence alignment of putative *TAM* orthologs from maize, compared with *AtTAM* and *SlTAM*.

```
cyc9            MAMRKENPVLNACQASNGRITRAQAAANRRSFGTFPSVPLPAKTERKQTAQGKAKRGSSY      60
AtTAM           ---------------------MSSSSRN-----LSQENPIP----RPNLAKTRTSL----      26
SlTAM           ---------------------MMTT-QN-----RRSSAVAK----RQAMAANSSGL----      25
cyc27           ---------------------MSCWAAGRRSSASAM-------------AE-NTGA----      21
cyc2            ---------------------MSTIAASRRSSSSSSSATAK----RPAIADGP-------      28
cyc6            ---------------------MSTHAASRRSS--SSSAVAK----RPAIAEGATKA----      29
                                           .                     *

cyc9            DNTSASVALSGPQPKRRTVLRDVTNLSNANSNKSFAAA----------P-----------      99
AtTAM           ----------RDVGNRRAPLGDITNQKNGSRNPSP------------SSTLVNCSNKIGQ      64
SlTAM           ----EHNQTGKLNAKKRPALSNISNNTTVSARNSVSH----------SSKLAPCTSKIVN      71
cyc27           ----RGAKEAVVTSGKRVALGNLTNVFRGSWGSGAAN-------SASDAKLSSTKP--DD      68
cyc2            ----GGPKATAAQAKKRVALGNITNVAARGGRASVGGSLGNVMAPTSSAKLNPT----VP      80
cyc6            ----AGPGPTAAQAKKRTALVNITNVAAPGARVAAVG---KVAPPVTGAKLNPATSG-AP      81
                               :*  * :::*    .   .

cyc9            -------------------------KLQTKPSLRTGRTVSK---------SKPCAKKIPK      125
AtTAM           SKKAP------------------------KPALSRNWNLGILDSGLPPKPNAKSNIIVPY      100
SlTAM           IKKNTSACNSNAVS---SGTAVLPASSSVRPS-S--KPVSIQRSDAV----VPKITVI--      119
cyc27           VKKGSYACL-RN-VNTERGSSRRLASEQFDQ------AVSLLQKNTSLP-SVPDVV----      115
cyc2            LKKPSLATSARSVSSSIRGSADKPASIKPAPPVARHGSATQRHNNVPPP-KVPTIADVPS      139
cyc6            LKKPSLAN-----ARTIRGSAAKSASIKPAPPVSRHDSSSEQKHNVL----VPTTVHVPS      132

cyc9            KPPPAGNGSALTNVLNIAE--E---TQAEKIL----AERVEPVLLLENRG-PLSLQNVER      175
AtTAM           EDTELLQSDDSLLCSSPALSLDASPTQSDPSISTHDSLTNHVVDYMVES-----------      149
SlTAM           --PVPA---TCSMDISPSHSDGSL-VSMDESMSNSDTVRSPEVEYIDDHELAAV-DSIEK      172
cyc27           ----------STGCSSPGLSQDCS-VSMEDAMSTCNSTESSDVEYLNNNDDPSMTSSSHW      164
cyc2            RAPALV---SCTGLVSPGRSGD-S-VSSDETMSTCDSMKSPDFEYVDNQDTSM-LASLQR      193
cyc6            RAPALV---PYSSFVSPGRSRD-S-VSTDETMSTCDSMKSPDFEYIDNGGCSM-LASLQR      186
                               . .       .. :  :    :     .  : :

cyc9            NRDSACHEVFFEE-RNL--RDKCEPSVSKNGDSYVLDIVDIDKDNGNPQMCASYVVEIYS      232
AtTAM           ------------------------TTDD-GNDDDDDEIVNIDSDLMDPQLCASFACDIYE      184
SlTAM           ---KACSTLYISEHVKAAAADICKRDVL-VDLESGDKIMNIDNNLVDPQLCATMACDIYK      228
cyc27           ---WASDRPHFSDSMDVAEFNW--RKHS-PN----------HYTYKDPRLSTTLACEIYE      208
cyc2            ---RTSEHLRISEDRDVEENKR--KKNA-VAPMEIDRICDVDSEYEDPQLCATLASDIYM      247
cyc6            ---RADEHLRTSEDRDVEENKW--KKNG-PAPMEIDSICEVDSNLEDPQLCAALASDIYM      240
                                                         .    :*::.:: . :**

cyc9            NLMASELMRRPSPNYMEGLQRDITKGMREILIDWLVEVSEEYKLVPDTLYLTVYLIDRFL      292
AtTAM           HLRVSEVNKRPALDYMERTQSSINASMRSILIDWLVEVAEEYRLSPETLYLAVNYVDRYL      244
SlTAM           HLRASEAKKRPSTDFMAKVQKDINPSMRAILIDWLVEVAEEYRLVPDTLHLTINYIDRYL      288
cyc27           SLREAETRKMPPTNFLETTQTDMSKTMRAILIDWLVEVTEEYRLVPETLYLTVNYIDRYL      268
cyc2            HLREAETKKRPSTDFMETIQKDVNPSMRAILIDWLVEVAEEYRLVPDTLYLTVNYIDRYL      307
cyc6            HLREAEMKKRPSTDFMKTIQKDVNPSMRAILIDWLVEVAEEYRLAPDTLYLTVNYIDRYL      300
                 *  :*  : *  :::   * .:.  ** *********:***:* *:**:*::  :**:*

cyc9            SRNYIERQRLQLVGITSMLVASKYEEICAPRVEEFCFITDNTYTKAEVLKMESQLLNDLG      352
AtTAM           TGNAINKQNLQLLGVTCMMIAAKYEEVCVPQVEDFCYITDNTYLRNELLEMESSVLNYLK      304
SlTAM           SGNLMDRQRLQLLGVACMMIASKYEEICAPQVEEFCYITDNTYFKEEVLQMESAVLNYLK      348
cyc27           SVKEISRHRLQLVGVACLLIAAKYEEICPPLVEELCYITDYSYTKEEVLQMEASVLNNLK      328
cyc2            SGNEISRQRLQLLGVACMLIAAKYEEICAPQVEEFCYITDNTYFRDEVLDMEASVLNYLK      367
cyc6            SGNEINRQRLQLLGVACMLIAAKYEEICAPQVEEFCYITDNTYFRDEVLEMEASVLNYLK      360
                : : :.::.***:*::.:::*:****:* * **::*:*** :* : *:*.**: :** *

cyc9            FNLSVPTTKTFLRRFLRAAQASRKTPSMTLGFLANYLAELTLTEYEFLKFLPSLVAASAV      412
AtTAM           FELTTPTAKCFLRRFLRAAQGRKEVPSLLSECLACYLTELSLLDYAMLRYAPSLVAASAV      364
SlTAM           FEMTAPTAKCFLRRFVRAAQGLNEVLSLQLEHLASYIAELSLLEYNMLCYAPSLIAASAI      408
cyc27           FEMTVPTAKCFLRRFVRVAQVPDKGSSLHLEFLVNYICELSLLEYSLLCYLPSMVAASSV      388
cyc2            FEMTAPTAKCFLRRFARAAQACDEDPALHLEFLANYIAELSLLEYSLLSYPPSLIAASAI      427
cyc6            FEMTAPTAKCFLRRFARSAQACDEDPALHLEFLANYIAELSLLEYSLLSYPPSLIAASAI      420
```

```
           *:::.**:* ***** * **   :  ::    *. *: **:* :* :* : **::***::

cyc9       FLARWTLDQSDLPWNQTLEHYTSYKCSDIQLCVCALRELQHNTSNCPLNAIREKYRHQKF     472
AtTAM      FLAQYTLHPSRKPWNATLEHYTSYRAKHMEACVKNLLQLCNEKLSSDVVAIRKKYSQHKY     424
SlTAM      FLAKYILLPSVKPWNSTLRHYTLYQPSDLRDCVLALHSLCCNNNNSSLPAVREKYSQHKY     468
cyc27      FLAKFILMPIKNPWNSSLSYYTRYTPSELRGCVRVLHRLFRLGPGSNLPAIREKYSQHKY     448
cyc2       FLARFILQPTKYPWNSTLAHYTQYKPSKLSECVKALHRLCSVGSGSNLPAIREKYSQHKY     487
cyc6       FLARFVLQPTKYPWNSTLAHYTQYKPSELSECVKTLHRLSSVGPGSNLPAIREKYSQHKY     480
           ***:: *     *** :* :** *  ..:  **  *  *     .. : *:*:** ::*:

cyc9       ECVANLT-SPEFPRSFFS----     489
AtTAM      KFAAKKLCPTSLPQELFL----     442
SlTAM      KFVAKKYCPPTVPVEFFQNISS     490
cyc27      KFVAKKYCPQSIPTKFFQDLTS     470
cyc2       KFVAKKQCPPQIPTEFFRDTTC     509
cyc6       KFVAKKQCPPQIPAEFFRDAAC     502
           : .*:      .* .:*
```

Identity matrix

| | cyc9 | AtTAM | SlTAM | cyc27 | cyc2 | cyc6 |
|---|---|---|---|---|---|---|
| 1: **cyc9** | 100.00 | | | | | |
| 2: **AtTAM** | 41.38 | 100.00 | | | | |
| 3: **SlTAM** | 41.57 | 51.17 | 100.00 | | | |
| 4: **cyc27** | 39.85 | 45.73 | 48.24 | 100.00 | | |
| 5: **cyc2** | 41.31 | 50.58 | 57.83 | 58.10 | 100.00 | |
| 6: **cyc6** | 40.95 | 49.42 | 56.78 | 56.52 | 84.48 | 100.00 |
